# Feedforward computational models of vision do not explain expert neural processing of visual Braille in the human visual system

**DOI:** 10.64898/2026.04.14.718353

**Authors:** F. Cerpelloni, O. Collignon, H. Op de Beeck

## Abstract

The human visual system, and in particular the Visual Word Form Area (VWFA), adapts to process letters and words, even when the stimuli do not share canonical script features, like Braille. Here we set-up to compare the organization of typical orthographic and peculiar visual scripts such as Braille in computational models. In a first experiment, we looked at how Braille letters are represented in an illiterate Convolutional Neural Network (AlexNet) and compared them to Latin alphabet and to Line Braille, a custom line-based script. We observed a predisposition of the network, pre-trained to perform object recognition, for line-based scripts. This finding suggests an initial advantage of line junctions over Braille in processing scripts likely based on typical visual computations applied to the visual world. In a second experiment, we trained two benchmark neural network architectures (AlexNet, CORnet Z) to classify words in the Latin script (literacy acquisition) and then in the Braille script (expertise acquisition). We modelled the processing of reading visual Braille and explored the networks representations at different layers. We observed clustering of features based on the visual properties of the scripts and not by the network’s expertise. Unlike human participants, the representations of linguistic categories do not converge to a model of the linguistic (orthographic, phonological, semantic) properties. Overall, the lack of alignment between the visual processing of the trained computational models and neural data recorded in expert humans suggests that the fundamental processing of reading cannot be fully explained by simple feed-forward visual processing of the script, but likely relies on additional mechanisms including interactive relations between the visual and linguistic systems.

## Introduction

In humans, the Visual Word Form Area (VWFA) has been found to show a preference for written scripts over other categories typically processed in the ventral occipito-temporal cortex (VOTC) (Cohen et al., 2000, 2002; Li et al., 2024). The first models of how VWFA acquires its selectivity for orthography emphasised the progressive integration of line junctions in the visual stream (Dehaene et al., 2005; Vinckier et al., 2007) building on a pre-existing visual hierarchy (Dehaene et al., 2005, 2015; Dehaene-Lambertz et al., 2018), and found that line vertices posed an advantage both behaviourally and in the brain in both object and word recognition (Szwed et al., 2009, 2011). This led to the hypothesis that line junctions and the visual system’s predisposition to process these features might be necessary to achieve fluent reading (Bola et al., 2017). However, other behavioural findings showed that once script novelty and training paradigm are equated, the advantage of line junctions in learning a new script is small and limited to the earlier stages of the learning phase (Cerpelloni et al., 2026), while the linguistic content of the stimuli learnt seemed to determine the learning curves. This last result is in line with an interactive account of VWFA (Price & Devlin, 2003, 2011) that stresses the role of VWFA in other tasks involving language (e.g. colour naming; Price & Devlin, 2003) and its interactions and connectivity with areas of the language network as the building block of script preference in those occipital regions (Price & Devlin, 2003, 2011; Saygin et al., 2016; S. Wang et al., 2022; X. Wang et al., 2018). Recent studies on the neural organization of VWFA showed that this regions implements representations of diverse linguistic properties (i.e. orthography, phonology, lexicality) in segregated neural populations (Planton et al., 2019; S. Wang et al., 2025) based on a diverse set of linguistic stimuli, as for example lip-reading (Van Audenhaege et al., 2025), auditory phonemes or words (Pattamadilok et al., 2019; Van Audenhaege et al., 2025) or for the reading of visual Braille (Cerpelloni et al., 2025).

To study the respective contributions of visual and linguistic properties in reading, one approach is to study them in isolation from one another, to identify the individual contributions of each to the whole process of visual interpretation of symbols and its relation to linguistic units. This is made possible by computational models of the visual system. The recent development of Deep Neural Networks (DNNs) made available in-silico models of the visual processing hierarchy, isolated from other cognitive functions that may contribute to object recognition (e.g. top-down connections with language areas, attention). Among a large corpus of computational studies focused on object recognition across categories, a few studies looked at the specific case of reading. First, Janini and colleagues tested a benchmark Convolutional Neural Network (AlexNet; Krizhevsky et al., 2017) on letter recognition and observed that training on naturalistic images (ImageNet dataset; Russakovsky et al., 2015), rather than specifically on letters, leads to letter identity representations that are better correlated with human behavioural data (reaction time in identifying different letters) (Janini et al., 2022). Such result supports the idea that the visual hierarchy of object recognition is co-opted by reading (Dehaene et al., 2005). Second, to achieve a synthetic model of VWFA, Hannagan, Agrawal, and Dehaene (Agrawal & Dehaene, 2024; Hannagan et al., 2021) employed a network architecture considered more biologically plausible (CORnet Z; Kubilius et al., 2018). The network was trained to resemble the acquisition of an alphabet in children, who are exposed to object prior to formal schooling (training on ImageNet) and then are introduced to words (training on ImageNet and word categories). In the Latin alphabet (French words), the authors observed ordinal and position coding of letters in a word. The extent of orthographies tested in isolation or in combination makes this model of reading a solid framework to study literacy acquisition and representations in deep neural networks. Nonetheless, the cases studied all rely on line-based scripts and are not generalized to non-canonical forms of visual reading like Braille, sign language, or lip reading, all found to be represented in VWFA similar to more canonical line-based orthographies (Cerpelloni et al., 2025; Van Audenhaege et al., 2025). It is unclear whether the correspondences between in-sillico models, human behaviour, and VWFA hold true in these cases, where the linguistic content is not conveyed through line-junctions.

Here, we tested two DNN architectures, previously used in studies on in-silico reading to represent humans behaviour and neural correlates, on the internal layer representations when presented with letters and words in the Latin and Braille alphabets, and compared the networks representations to human results. Behaviourally, the comparison of participants who learnt Braille or a script based on line-junctions shows an early advantage of line-junctions that is however quickly compensated by the linguistic content mapped between the native and novel alphabet learnt, regardless of the visual features (Cerpelloni et al., 2026). At the neural level, VWFA shows similar univariate responses to visual Braille and to Latin-based scripts, in addition to multivariate patterns common across scripts and correlated with a cumulative linguistic content (Cerpelloni et al., 2025). We presented stimuli with different visual (presence or absence of line junctions) but shared linguistic features (phonology, semantic) to word-recognition models and tested whether the visual processing hierarchy (without linguistic information) is sufficient to process the statistical regularities of the letters and words. Alternatively, a failure of these models in replicating these findings would suggest that visual processing by itself cannot trivially explain the human performances and that the interactions between visual and language networks might be needed to explain the findings in human participants. In a first experiment (Figure 1A), we explored how an illiterate visual network (i.e. not explicitly trained on alphabetic stimuli) represents letters from the Latin alphabet, from Braille, and from a custom script with Braille dots joined into lines (Line Braille). Similarly to the advantage in learning a line-based script found at the beginning of training, we observed higher similarities between Line Braille and Latin alphabet, both showing low similarity with Braille, despite our procedure to base the Line Braille upon the Braille structure. In a second experiment, we looked at the representations of full words, whether known or not by the network. We fine-tuned previously trained networks using Braille or Line Braille, simulating the behavioural training experiment (Cerpelloni et al., 2026). We found an advantage in learning a line-based script over Braille, an effect that was in the same direction as the behavioural findings but much larger in magnitude and duration for the networks. After training, we tested (Figure 1C) the networks trained on Braille (Braille-expert) and trained solely on the Latin alphabet (Braille-naïve) on the representations of stimuli in Latin and Braille alphabets which varied in the linguistic properties (Real Words, Pseudo Words, Non Words, Fake Script), to compare the network and the brain organizations. Overall, network expertise impacted the clustering of representations only marginally (i.e. the difference between two categories over the difference within them). The multivariate relationship between networks representations at the output layers showed similar representation of orthographies rather than correlations based on the training or the architecture. Differently from what is found with human participants, these representations did not correlate with a linguistic model organization. These findings identify a gap in the current state of computational models aimed at reading, showing that the word recognition performed by computational models of vision is not sufficient to explain the representation of the same stimuli in the human mind and brain and highlighting instead the need to integrate visual processing with linguistic one.

**Figure 1.**
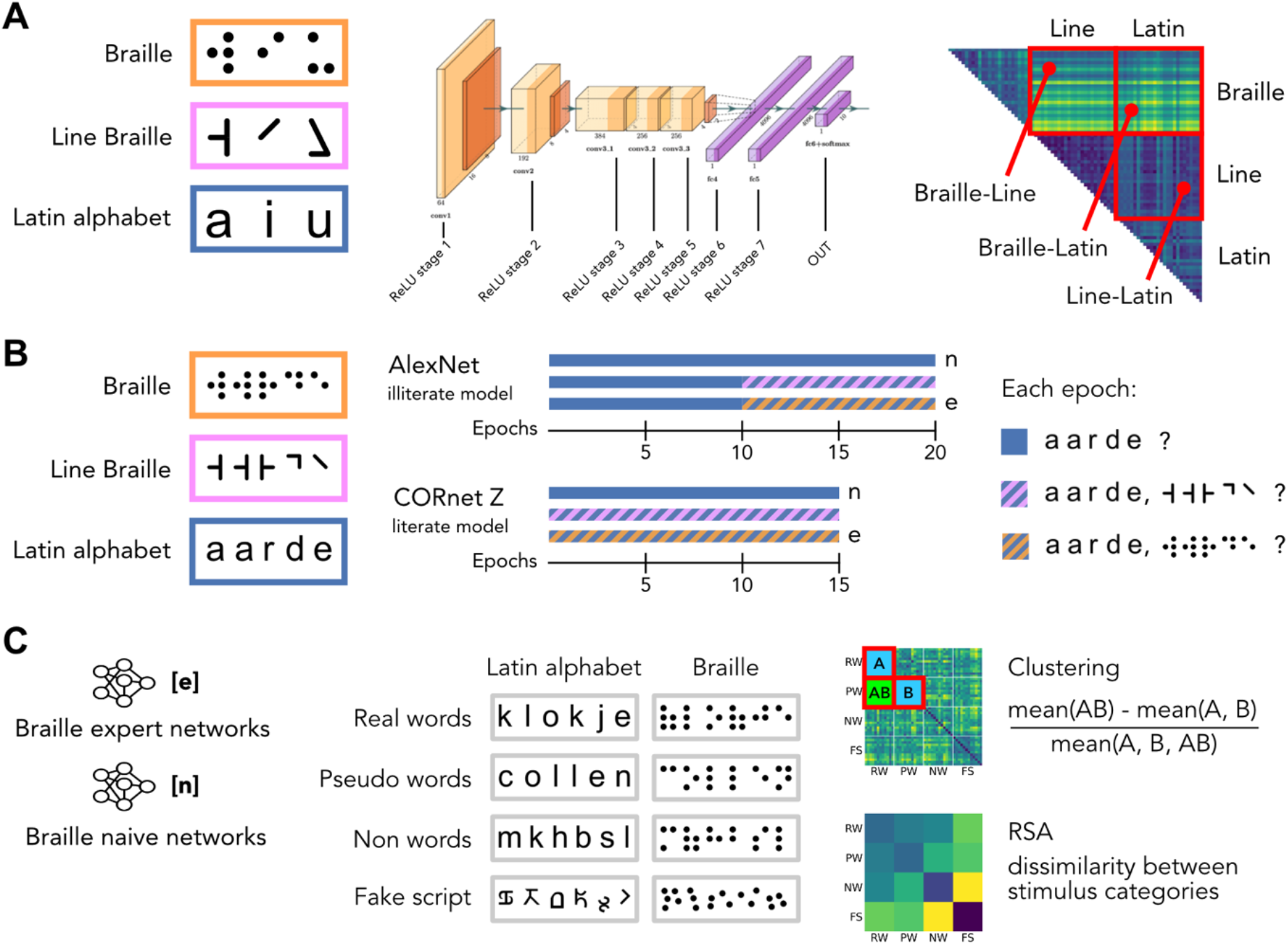
Experiments design. (A) Experiment 1, representations of letters in illiterate AlexNet. Letters in the Latin alphabet (Arial font), Braille, and Line Braille were presented to a AlexNet instance pre-trained on ImageNet but not explicitly on letters. The activations in response to each letter at different processing stage was used to compute the Euclidean distance and to assess the differences between alphabets. (B,C) Experiment 2, representations of words and expertise acquisition. (B) Word stimuli from the three alphabets were presented to AlexNet and CORnet Z models. AlexNet was first trained on the Latin alphabet alone and then Braille or Line Braille were introduced alongside the Latin alphabet. CORnet Z, from previous studies where it was trained on the Latin alphabet, was directly trained on the specific dataset of this experiment in all three datasets: Latin alphabet alone, Latin alphabet + Line Braille, Latin alphabet + Braille. At the end of each epoch, the networks were tested on the recognition of a subset of the words trained, in each alphabet trained. (C) Networks trained solely on Latin alphabet (n; naïve) and on Latin alphabet + Braille (e; expert) were tested on a set of stimuli with different linguistic (orthographic, phonological, semantic) properties in both Braille and Latin alphabets. The activations were used to measure clustering and dissimilarity between categories at different processing stages of both AlexNet and CORnet Z models.

## Methods

### Stimuli

To explore the role of line junctions on the script representations, we compared three scripts: Braille, Latin-based, and a custom ‘Line Braille’ script made of lines previously used to investigate behavioural learning (Cerpelloni et al., 2026). All stimuli were controlled for the size of the letters.

In experiment 1, we explored the representations of letters in a AlexNet model considered “illiterate”, not explicitly trained on letters or words. For each script, we selected the 26 letters of the Latin alphabet and created images for each of the three scripts, manipulating the thickness and size of the letters to introduce variability in the stimulus set. Thickness variations consisted in adding or removing pixels from the perimeter of the letter, resulting in bolder or thinner features. We chose five variations consisting of -6, -3, 0 (original image), +3, +6 pixels layers. Similarly, size variations consisted in five levels of increase of the letter dimensions, resulting in letters that were -30%, -15%, 0%, +15%, +30% the size of the original letters. All final images were 500×500 pixels, with the letter presented in black at the centre of a white background. In experiment 2, we assessed the contributions of the visual features to reading. We referred to the methods used by Agrawal and Dehaene (2024) and constructed different datasets to train and test the word recognition models (see *DNN architectures and training*), representing the literacy and expertise of the different groups who underwent Braille or Line Braille learning: (1) a dataset of words solely in the Latin-based alphabet (LT) to mimic participants naïve to either Braille or Line Braille; (2) a dataset of words in the Latin-based and Braille alphabets (LTBR), to mimic individuals who underwent Braille behavioural training (Cerpelloni et al., 2026) and those with extensive expertise leading to VWFA accommodation (Cerpelloni et al., 2025); (3) a dataset of words in the Latin-based and Line Braille alphabets (LTLN), to mimic the individuals who learnt Line Braille in the behavioural training. All datasets included words in the Latin-based alphabet to reflect the training performed by human participants, who learnt the new script without losing exposure to Latin-based Dutch. All datasets contained the same one thousand Dutch words from the Dutch Lexicon Project (Brysbaert et al., 2016), but varied in the number of fonts present in the set. All words selected were frequent (top 50% of the dataset) and balanced in length, component (i.e. piece of sentence), and orthographic neighbours. A dataset comprised of either 1100 or 1375 stimuli per word, depending on the training set: all datasets contained four Latin-based alphabet fonts (Arial, Times New Roman, American typewriter, Futura), while the datasets with novel alphabets added an additional font of Braille or Line Braille. Each word within a font was varied in size (-30%, -15%, 0%, +15%, +30% of the original size), and shifted on the x-(-50, -40, -30, -20, -10, 0, 10, 20, 30, 40, 50 pixels shifts) and y-axes (-40, -20, 0, 20, 40 pixels shifts) from the center of the image. To assess the accuracy of learning at each epoch, we presented a subset of the stimuli used in the training to the networks. We selected 20 words balancing the number of letters in the stimulus and the other linguistic properties. In the Latin alphabet, we selected words in the Arial font and extracted all their stimuli variations. In the Braille and Line Braille scripts, we selected the same words in all their variations. To test the within-layer representations of stimuli with different linguistic (i.e. semantic, phonological, orthographic) content, we constructed a test set based on stimuli from a previous fMRI experiment focused on the neural organization in VWFA (Cerpelloni et al., 2025). Following this method, the stimulus set included twelve stimuli for each of the linguistic categories: (1) Real Words (RW), selected from the training set classes (i.e. semantic meaning) and thus made of high-frequency bigrams and trigrams and known letters; (2) Pseudo Words (PW), built using unipseudo (New et al., 2024) from a subset of the training set words and including only strings with high-frequency and familiar letters but without an output class associated; (3) Non Words (NW), strings of consonants composed of low-frequency bigrams and trigrams; (4) Fake Script stimuli (FS) obtained by transcribing the Non Words into a unfamiliar script made of re-arranged line-junctions. We adapted the Real Words and Pseudo Words sets (related to each other) to Dutch and to the specific requirements of this study and used the same stimuli of the fMRI experiment for the sets of Non Words and Fake Script, as the strings of letters did not contain frequent bi- or tri-grams. All stimuli were presented in both Latin-based (Arial) and Braille alphabets, and underwent the same size variations and axes shifts of the training sets.

### Deep Neural Networks (DNNs)

In experiment 1, we presented these stimuli to an “illiterate” AlexNet, pre-trained on ImageNet to recognize objects but not explicitly trained on letters or words. We then extracted the responses at each layer (each ReLU stage and output) to the representations of single-letter stimuli in the three scripts, following the methods of Janini and colleagues (2022). In experiment 2, we implemented two neural network architectures from previous studies: AlexNet (Krizhevsky et al., 2017) and CORnet Z (Kubilius et al., 2018) previously trained to recognize words in the French alphabet (Agrawal & Dehaene, 2024; Hannagan et al., 2021).

For AlexNet, we implemented instances (N = 5) of the network pre-trained on ImageNet (Russakovsky et al., 2015), referred to as “illiterate”, and reset the output classes while maintaining the weights of the other layers intact. Since AlexNet models were “illiterate”, we first trained the models on the Latin alphabet stimuli set (training phase of literacy acquisition; learning rate = 1e-4; batch size = 100; training set = 70% of the full dataset). After 10 epochs, the networks reached a classification accuracy above 95% and we introduced expertise for a novel script. We obtained three duplicates of the same network instance and each copy was further trained for 10 additional epochs on one of the three datasets (phase of expertise acquisition): introducing Braille (further trained on the Lain-based and Braille alphabets; N = 5), Line Braille (further trained on the Lain-based and Braille alphabets; N = 5), or continuing with the Latin alphabet training (further trained on the Latin-alone dataset; N = 5).

We additionally implemented a CORnet Z model previously trained on ImageNet and then on ImageNet and words (Agrawal & Dehaene, 2024; Hannagan et al., 2021), considered a synthetic model of VWFA. We started from the French-literate network presented in Agrawal and Dehaene (2024) and adapted the number of output units to match the one thousand words we selected for the training datasets. CORnet models, already trained on a dataset of words and therefore considered literate, were trained in three duplicate instances on one of the three words datasets, solely on the Latin alphabet or adding either braille or Line Braille. Each instance was trained for 15 epochs, with the same hyper-parameters used for AlexNet (learning rate = 1e-4; batch size = 100; training set = 70% of the full dataset). We extended the training epochs in CORnet to include learning of the novel categories.

At each epoch, we extracted and saved the weights of each network instance. To assess the accuracy of the training at each epoch, we presented the network (with frozen nodes to prevent further learning) with stimuli coming from the training dataset and obtained the output classification. After training, we tested the networks trained solely on Latin alphabet and with the addition of Braille against our test dataset of linguistic conditions in both scripts. In both AlexNet and CORnet networks, we extracted the features activations at different layers (AlexNet: ReLU stages of each layer, and output; CORnet: output layer of each CORBlock, linear and output layers of decoder) for each stimulus image presented to the network.

### Statistical analyses

To measure the dissimilarity between letters identities in Experiment 1, we computed at each ReLU stage and output layer the Euclidean distance between stimuli and compared the average response across variations to one stimulus (one letter in one alphabet) to all the individual variations of another stimulus. We then averaged the distances and repeated the same computation across all letters in all alphabets, following the methods described by Janini and colleagues (Janini et al., 2022). This process resulted in a Representational Dissimilarity Matrix (RDM) indicating the Euclidean distance between each letter in each alphabet to all the other letters and alphabets. We then computed mean distances between each alphabet and compared them using a permutation test (10,000 label shuffles of the shared category), correcting for multiple comparisons using Bonferroni correction. In experiment 2, we assessed the different learning curves associated to the Latin and Braille, and Latin and Line Braille alphabets datasets. In particular, we compared the responses of the networks to a subset of training words at each epoch of the expertise phase through a repeated measures ANOVA (rmANOVA) for each network (2 scripts * 10 epochs in AlexNet; 2 scripts * 15 epochs in CORnet Z). To test the emergence of representations due to expertise, we first replicated the analyses of Agrawal and Dehaene (Agrawal & Dehaene, 2024) and computed the mean dissimilarity (1-correlation) between the activations of each layer of the network for each stimulus, and then averaged the values to obtain a measure for each layer. We also computed RDMs of the Euclidean distance for each word identity in each layer and network, separating each script (Braille and Latin alphabet), using the same methods used for the letter identities (Janini et al., 2022). We then computed a measure of the clustering of the representations across the four different linguistic conditions (Real Words, Pseudo Words, Non Words, Fake Script). We defined clustering as the average dissimilarity in comparisons between conditions minus the dissimilarity within conditions, divided by the overall average dissimilarity of representations (see Figure 1C). We then averaged the clustering of each condition to obtain a measure of the overall specialization of one layer. To explore effects of expertise in the clustering across scripts, we performed rmANOVAs on the average clustering (2 scripts * 2 network expertises * 8 layers in AlexNet or 6 layers in CORnet Z). To explore the relationships between stimuli representations of the different scripts and expertise, we averaged the RDMs at the output layer of each network to obtain representations of the linguistic categories. Within a network architecture, we correlated the representations of different scripts in different network expertises among each other, and also with a theoretical model of organization of the linguistic categories, from a previous study using multivariate fMRI (Cerpelloni et al., 2025). The model was based on the number of linguistic properties (i.e. orthography, phonology, semantics) of each stimulus condition, with the dissimilarity given by the different number of properties between stimuli classes (e.g. the difference between Real Words, possessing orthographic, phonological, semantic properties, and Pseudo Words, possessing orthographic and phonological, but not semantic properties, would be one as there is one level difference). Given the small sample of network instances, we followed the same method and computed non-parametric statistics by performing bootstrapping (10000 samples) to obtain a null distribution of the correlations between matrices.

### Data and Code availability

All the in-house scripts coded to select and create the stimuli, as well as to train and test the networks, are publicly available on GitHub: https://github.com/fcerpe/VBS_data. All the weights of the network and the activations extracted in response to the test set is available upon request through GIN G-Node: https://gin.g-node.org/fcerpe/VBS_weights and https://gin.g-node.org/fcerpe/VBS_activations.

## Results

### Letter processing depends on the presence of line junctions

We presented letters in the Latin alphabet (Arial font), in Braille, and in a custom Line Braille script to an instance of AlexNet pre-trained on ImageNet but illiterate (i.e. not directly trained on written material) and measured the network activations at different processing stages (Figure 2). We observed an increase in the dissimilarities between scripts, and a progressive bias for lines over Braille emerging from the mid-layers of the network (ReLU stage 3). In particular, in the first feature layer (layer 1; Braille/Line – Braille/Latin distance: 25.1, p_FDR_ = 0.36; Braille/Line – Line/Latin distance: 50.6, p_FDR_ = 0.06; Braille/Latin – Line/Latin distance: 25.5, p_FDR_ = 0.29) we did not observe significant differences in the distances between scripts, meaning that the alphabets are processed without line-junctions biases but on lower-level properties. In contrast, the distances between Braille and line-based scripts are significantly greater than the distances between Line Braille and Latin alphabets In mid-level features layers (layer 3: Braille/Line – Line/Latin distance: 143.3, p_FDR_ < 0.001; Braille/Latin – Line/Latin distance: 98.6, p_FDR_ = 0.003. Layer 4: Braille/Line – Line/Latin distance: 68.2, p_FDR_ < 0.001; Braille/Latin – Line/Latin distance: 45, p_FDR_ = 0.03. Layer 5: Braille/Line – Line/Latin distance: 29.9, p_FDR_ = 0.006; Braille/Latin – Line/Latin distance: 30.9, p_FDR_ = 0.003) and in the fully-connected layers (layers 7: Braille/Line – Line/Latin distance: 1.83, p_FDR_ = 0.023; Braille/Latin – Line/Latin distance: 1.49, p_FDR_ = 0.046. Output: Braille/Line – Line/Latin distance: 3.01, p_FDR_ < 0.001; Braille/Latin – Line/Latin distance: 1.86, p_FDR_ = 0.021). Line Braille and Latin alphabet present significant distances only in the second layer, where major differences between all scripts emerge (all p_FDR_ < 0.001). This result shows a preferential processing of line junctions in a network trained solely on natural objects, and reveal the emergence (after the first layer) of a distinct visual representation for a script that does not share those basic features.

**Figure 2.**
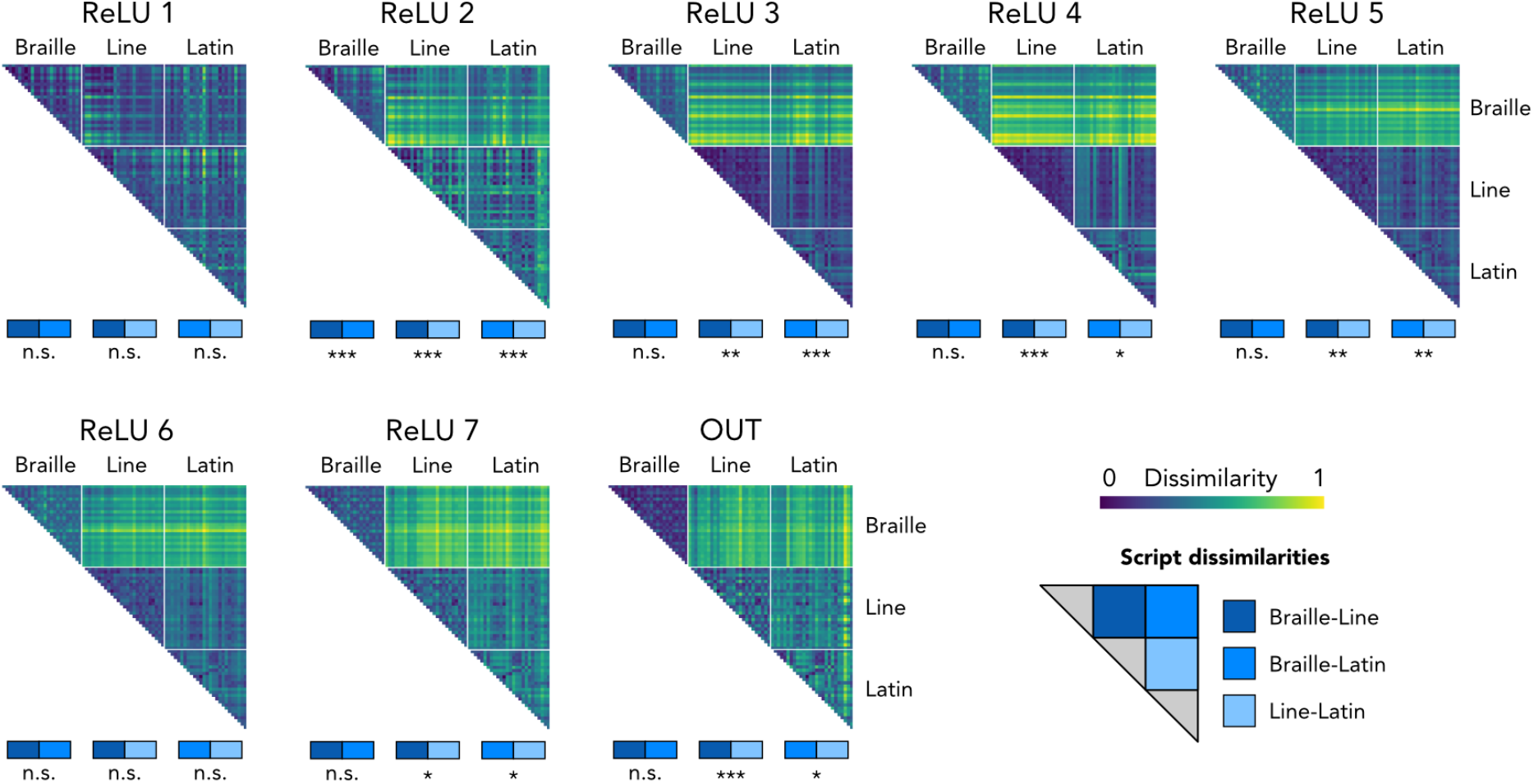
Illiterate AlexNet representations of letters in different scripts. Letters in Braille, in Line Braille, in Latin alphabet were presented to the network (illiterate, not explicitly trained on letters). The network activations extracted at given stages (ReLU stages and output) for all the individual letters in their size and thickness variations were compared across scripts by computing the Euclidean distance between stimuli, obtaining a measure of the letter identity. The Representational Dissimilarity Matrices (RDMs) show different processing stages for all the letters of Braille, Line Braille, Latin alphabet respectively. From early layers, a trend of statistical differences between Braille and other alphabets emerges across the visual hierarchy. In all the plots, stars represent significance (* p ≤ 0.05, ** p ≤ 0.01, *** p ≤ 0.001).

### Expertise acquisition shows benefits of line-based scripts over Braille

We trained AlexNet to first classify Dutch words presented in four Latin-based fonts (literacy acquisition) and then trained further solely on the Latin alphabet or introduced either Braille or Line Braille (joining the dots to form line junctions) alongside the Latin alphabet fonts, mapping the new script on the same output units (expertise acquisition). We extended the training of AlexNet in the novel and in the Latin scripts well beyond the convergence of the learning curves of Latin and Line Braille accuracy. We observed a drop of performance in both expertise conditions, stronger for Braille than for the line-based script (Figure 3A). Overall, across the ten epochs of expertise acquisition, we observed a main effect of epoch (F_(9, 72)_ = 2414, p < 0.001, η2_p_ = 0.99), indicating that both networks improve in the accuracy over training, and of script (F_(1, 8)_ = 19784, p < 0.001, η2_p_ = 0.99), which highlights different learning curves between Braille and Line Braille, given by the (lack of) line junctions. Additionally, we also found an interaction between script and epoch (F_(9, 72)_ = 306, p < 0.001, η2_p_ = 0.97). In the case of CORnet Z training (Figure 3B), starting from literate weights, we trained in parallel different instances on the three datasets (only Latin alphabet, or adding Braille or Line Braille to the Latin alphabet words), again for a longer number of epochs than needed for the convergence of accuracy ratings between Latin alphabet and Line Braille. We observed a temporal delay in the classification accuracy for novel scripts compared to the training on the Dutch words in Latin-based fonts. Moreover, while the expertise in Line Braille is compensated at epoch 10, the expertise in Braille showed a more marked delay. Indeed, with an average classification accuracy at epoch 10 much lower than the line-based trained networks (Braille: 0.5978, Latin alphabet alone: 0.9689; Line Braille expertise: 0.9435). A rmANOVA on epochs * novel scripts found main effects of epoch (F_(14, 112)_ = 418.3, p < 0.001, η2_p_ = 0.98) and script (F_(1, 8)_ = 162.4, p < 0.001, η2_p_ = 0.95), in addition to an interaction effect (F_(14, 112)_ = 69.38, p < 0.001, η2_p_ = 0.89). Overall, the expertise acquisition seems to align across network architecture in showing an advantage in introducing a script-based on lines rather than one void of those features. This strong bias for line-based alphabets diverges from the behavioural learning observed in humans (Cerpelloni et al., 2026), where the advantage posed by line-junctions was limited both in the duration and the magnitude of the effect.

**Figure 3.**
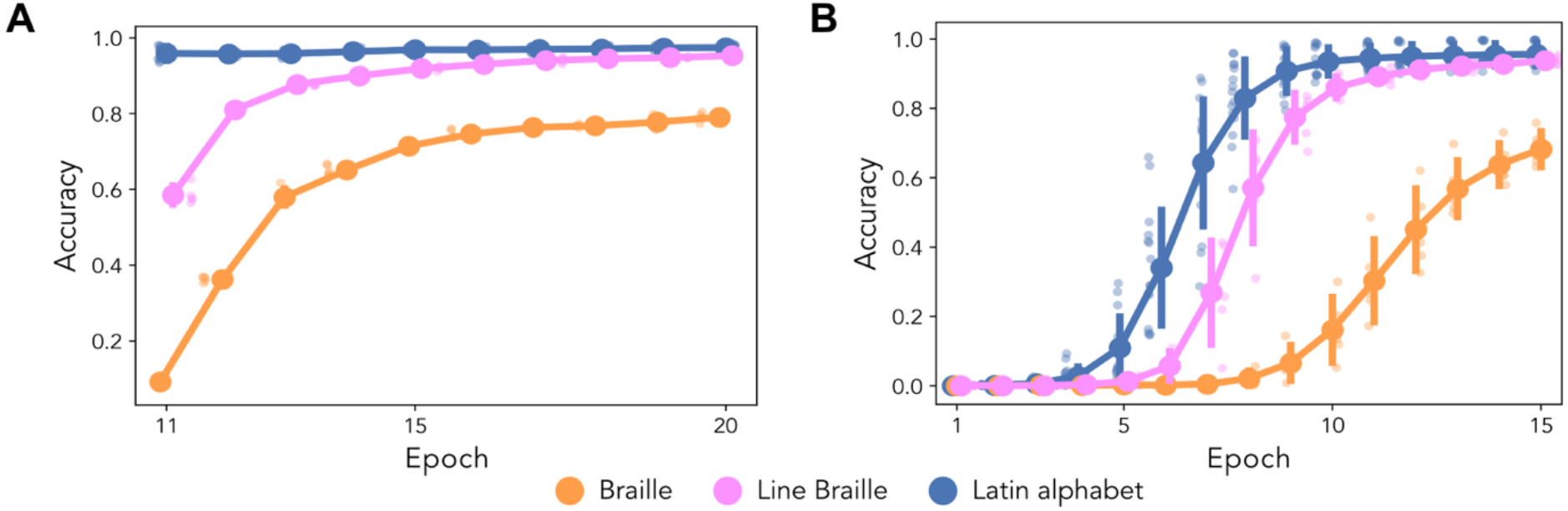
training results for the network architectures. (A) AlexNet results. We measured the classification accuracy of each network to a subset of trained words in their expertise script. The difference in classification of Braille words is impaired at the beginning of the expertise acquisition compared to Line Braille remains significant even when the performance of Line Braille matches the Latin alphabet. (B) CORnet Z results. Starting from pre-trained weights from Agrawal and Dehaene (2024), we further trained five instances of the network on each dataset of our experiment. Braille classifications show a delayed learning maintained at the end of the training regime. In both figures small dots represent single instances, vertical bars represent standard deviation (SD).

### Expertise acquisition does not influence word representations

We extracted the responses of both network architectures trained solely on Latin alphabet stimuli (naïve networks) and trained on Latin and Braille alphabets (expert networks) to stimuli that represented a gradient of linguistic (i.e. orthographic, phonological, semantic) properties. We first tested whether and at which layer the networks show an increase in mean dissimilarity, an indicator of improvement in the representations and therefore of expertise as a consequence of learning. In AlexNet (Figure 4A), there is a strong increase in the mean dissimilarity from layer 5, with trained alphabets showing a similar trend and the Braille alphabet in the naïve networks showing limited increase. In the CORnet models, we observed an increase in the mean dissimilarity starting from the layer IT, with higher mean dissimilarity for the Latin alphabet compared to Braille, regardless the network’s training. Only at the output layer we observed a difference between trained and untrained Braille representations. These results are already at odds with the result of Agrawal and Dehaene (2024), who showed an increase in the mean dissimilarity for trained alphabets from layer V4 (Figure 4B). We then tested whether expertise would result in a higher clustering of stimuli that belong to the same category (e.g., putting Pseudo-words apart from Non-words). We computed, for each network and layer, a measure of clustering (average dissimilarity in comparisons between conditions minus the dissimilarity within conditions, divided by the overall average dissimilarity) based on the dissimilarity of the activations in response to the test stimulus images. Using clustering, we performed a repeated measures ANOVA (2 scripts – Braille and Latin alphabet * 2 network trainings – expert and naïve to Braille * 6 or 8 layers, depending on the network). Overall, statistical analyses show very similar results in the two networks (Figures 4C and 4D), but with different directionality. More specifically, in AlexNet models (Figure 4C), the clustering of representations increases in association to the complexity of the processing stage (main effect of layer: F_(7,56)_ = 1626, p < 0.001), and a higher clustering is found in the Latin-based alphabet than in Braille (main effect of script: F_(1,8)_ = 3848, p < 0.001). However, whether the network was explicitly trained to associate Braille words to Latin-based categories does not play a role in how the network represents the different scripts (main effect of network’s expertise: F_(1,8)_ = 3.05, p = 0.12). We also found significant interactions between expertise and all the other variables (expertise * layer: F_(7,56)_ = 30.53, p < 0.001; expertise * script: F_(1,8)_ = 35.97, p < 0.001; layer * script: F_(7,56)_ = 408.77, p < 0.001; expertise * layer * script: F_(7,56)_ = 109.36, p < 0.001), driven by an increase in clustering of Braille stimuli in the fully-connected layers (layers 6 and 7, and output) of naïve AlexNet. This result can be explained by overall low dissimilarities between categories (Figure 4A), increasing the sensitivity of the clustering measure which then reflects small variations between categories rather than similar processing of different categories (i.e. linguistic) categories (expert, Braille mean dissimilarity before output layer: 0.27; naïve, Braille mean dissimilarity before output layer: 0.08). In CORnet Z models (Figure 4D), clustering is mainly determined by the processing stage (main effect of layer: F_(5,40)_ = 954.95, p < 0.001) and by the script processed (main effect of script: F_(1,8)_ = 798.82, p < 0.001), but not by the network expertise with Braille (main effect of expertise: F_(1,8)_ = 1.66, p = 0.23). Moreover, significant interactions between expertise, layer and script (expertise * layer: F_(5,40)_ = 41.72, p < 0.001; expertise * script: F_(1,8)_ = 211.47, p < 0.001; layer * script: F_(5,40)_ = 39.22, p < 0.001; expertise * layer *script: 16.83, p < 0.001) are present. We can observe visually in Figures 4C and 4D that the way the clustering for Braille evolves across layers in the expert and naïve network is opposite in CORnet (orange line goes lower than red line towards later layers) relative to AlexNet (orange line goes higher than red line).

**Figure 4.**
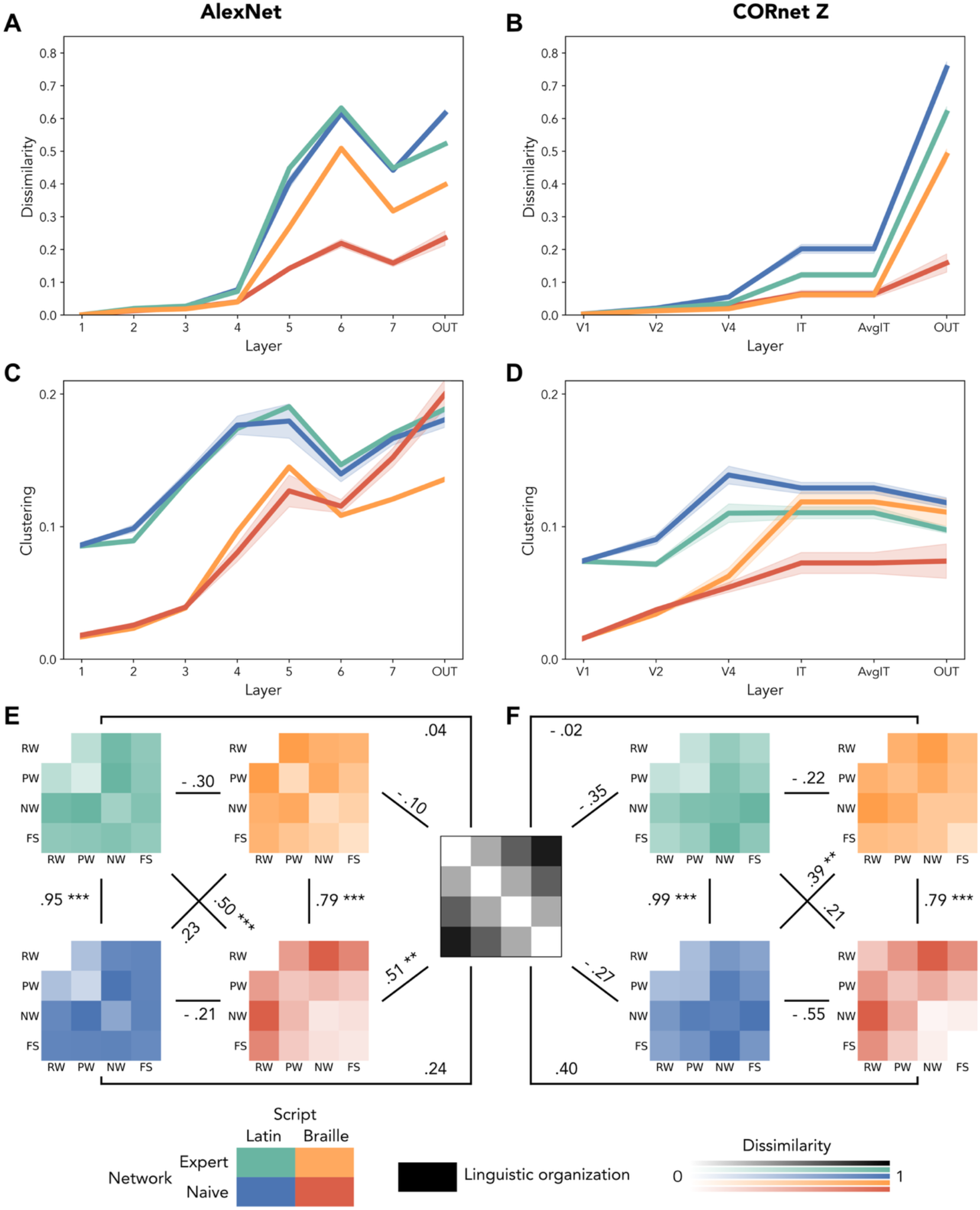
Network representations of words. (A, B) Mean dissimilarity across stimuli. AlexNet (A) and CORnet (B) both show an increase in the mean dissimilarity between test stimuli of a script in the middle to last layers, more marked for scripts that have been presented during training compared to novel scripts. (C, D) Clustering of test categories across layers. (C) AlexNet models show higher clustering (more dissimilarity between categories than within categories) for the Latin alphabet than for Braille, and for Braille script in the networks not trained on it. (D) CORnet Z models show an effect of expertise for Braille stimuli, with comparable levels of clustering to the Latin alphabet (extensively trained in all conditions) and Braille in the last layers. In A and B, the shaded area indicates 95% confidence interval. (E, F) Representational Dissimilarity matrices (RDMs) of the dissimilarities averaged across categories and of the model of linguistic organization (black, center). (E) AlexNet presents mostly correlations between Latin alphabet and between Braille representations across networks, in addition to a significant correlation between the untrained Braille representations and the linguistic model. (F) CORnet Z presents a similar pattern of correlations between network trainings, with no correlations with the model. In neither case, the networks represent the trained alphabets according to their linguistic properties. Stars represent significant correlations In all the plots, stars represent significance (* pFDR ≤ 0.05, ** pFDR ≤ 0.01, *** pFDR ≤ 0.001). x

Next, we investigated the amount of dissimilarity between categories and whether these category dissimilarities showed patterns as found before in humans (Cerpelloni et al., 2025). In the human brain, we found a graded pattern with e.g. less difference between Words and Pseudo-words than between Words and fake script. In addition, in experts these patterns were very similar between scripts. We focused on the dissimilarities between categories at the output layer of each network. In both networks, the output representations of linguistic categories do not show clear correlation patterns as shown in human results (Figures 4E and 4F). AlexNet (Figure 4E) presented significant correlations within scripts (expert, Latin – naïve, Latin: r = 0.95, p_FDR_ < 0.001; expert, Braille – naïve, Braille: r = 0.79, p_FDR_ < 0.001), and between Latin alphabet in the expert network and Braille alphabet in the naïve network (expert, Latin – naïve, Braille: r = 0.50, p_FDR_ = 0.0002). All the other correlations within and between networks (all p_FDR_ > 0.08) were not significant. Thus, there was no correlation between scripts in the expert network, contrary to the observation in humans. Similarly, in CORnet models (Figure 4F) we observed significant within-script correlations (expert, Latin – naïve, Latin: r = 0.99, p_FDR_ < 0.001; expert, Braille – naïve, Braille: r = 0.89, p_FDR_ < 0.001) and a significant correlation between Braille in the expert network and Latin alphabet in the naïve network (expert, Braille – naïve, Latin: r = 0.39, p_FDR_ = 0.004). All other correlation are not significant (all p_FDR_ > 0.09). Moreover, neither network architecture shows a clear trend of correlations with the theoretical linguistic model as found in human participants (Cerpelloni et al., 2025). While in CORnet Z models, no correlation is significant (expert, Latin – model: r = -0.36, p_FDR_ = 0.96; expert, Braille – model: r = -0.02, p_FDR_ = 0.96; naïve, Latin – model: r = -0.27, p_FDR_ = 0.96; naïve, Braille – model: r = 0.40, p_FDR_ = 0.083), in AlexNet, curiously, we only found a significant correlation between the Braille representations in the naïve AlexNet models and the theoretical RDM (expert, Latin – model: r = 0.04, p_FDR_ = 0.57; expert, Braille – model: r = -0.09, p_FDR_ = 0.68; naïve, Latin – model: r = 0.24, p_FDR_ = 0.22; naïve, Braille – model: r = 0.51, p_FDR_ = 0.02). While this result is unexpected and unexplained, it is clear that the representations formed by the networks do not represent the ones found in humans.

## Discussion

We explored the visual processing of Braille letters and words using computational models previously employed in word and letter recognition tasks (Agrawal & Dehaene, 2024; Hannagan et al., 2021; Janini et al., 2022). We first presented letters in the Latin alphabet (Arial font), in Braille, and in a custom script based on line junctions (Line Braille) to an instance of AlexNet, pre-trained on ImageNet. We observed that the absence of line junctions, as it is the case in Braille, led to higher dissimilarity of the letter representations, and that the presence of line junctions in Line Braille led to closer representations to Latin alphabet, also composed of line junctions, rather than Braille, a script sharing the same overall structure. We then trained AlexNet first to process Latin alphabet words and then also Line Braille and Braille words, and literate CORnet Z models to process either Latin alphabet words or Line Braille and Braille words. We observed a systematic bias for line-based scripts (Latin alphabet and Line Braille) compared to Braille, for who learning was significantly impaired throughout learning. We tested the Braille expert and Braille naïve (trained solely on the Latin alphabet) networks on stimuli with different linguistic statistics in both Latin alphabet and Braille. We found incongruent results between architectures in the clustering of representations, with different trends for trained and untrained Braille stimuli, both showing effects that underlying lack of expertise-induced differences. CORnet Z showed a significant interaction between expertise and script that can hint at an expert processing of Braille in the later stages (i.e. IT). In both networks, the main correlations between network’s representations were within scripts, while no correlation was present between a theoretical linguistic organization of stimuli and the representations of any trained alphabet, showing an overall discrepancy with how expertise changes representations in humans.

Letter processing in AlexNet naïve to orthography showed marked differences between Braille, based on dots, and line-based scripts, Latin alphabet and Line Braille. This result is particularly interesting first because Line Braille is derived from Braille’s same structure represented through lines instead of dot arrangements and would be expected to show more similarity to Braille than to the Latin alphabet. Instead, from the second layer onward line-based scripts are clustered together mostly without significant differences between them, indicating that the computational visual hierarchy tunes to line junctions (Agrawal & Dehaene, 2024; Hannagan et al., 2021; Janini et al., 2022). Experiment 1 shows that Braille is very much an outlier script for a visual network trained in general object recognition, strengthening the viewpoint that Braille challenges typical visual processing. Moreover, this result complements the behavioural data on the early learning of Braille (Cerpelloni et al., 2026) and adds to the significance of the behavioural learning of naïve participants, who show comparable results in learning either Braille or Line Braille after as little as a few hours of training.

Next we tested how artificial networks learn these scripts. To our knowledge, there are currently no computational studies directly comparing the acquisition of different orthographies. In this context, the only direct comparisons can be done with human studies. Cerpelloni, Collignon, Op de Beeck previously found that learning Braille or Line Braille results in a small initial advantage for Line Braille that is compensated in learning accuracy by the fourth consecutive day of training. In visual artificial neural networks we observed instead a much stronger bias for line-junctions, with no convergence in the learning accuracy at the end of the training epochs. While it remains possible that sufficient training of the networks would have resulted in a convergence of the scripts, the short number of hours spent by the human participants (approximately 4 hours each) is well reflected in the minimum ten epochs of network training. We trained the networks well beyond the point where the performance for Line Braille reached the asymptotic level and converged with the Latin script, and still the performance for Braille lagged. A longer training, with a possible convergence of scripts, would not hinder such divergence between the small and temporary delay in learning in humans and the large and long-term difference in computational models.

Agrawal and Dehaene (2024) tested the expertise of a network by comparing the activations of their models trained on either Telugu or Malayalam using bigrams from both scripts and found that the mean dissimilarity increased from the layer associated to area V4 only when the script matched the network expertise. Similarly, we found increasing dissimilarity in AlexNet from mid-layers while in CORnet only from layer IT and with lower dissimilarity associated to Braille, even if trained. We deepened the analysis by looking at clustering (dissimilarity between categories over the overall dissimilarity) and found the Latin alphabet representations to increase with the hierarchical processing stages. However, clustering was mainly determined by the script processed and only marginally by the expertise of the network, Braille in fact did not show a clear-cut effect of expertise. Moreover, the networks diverge in the clustering patterns of Braille stimuli. This divergence of the models can be partially explained by the low dissimilarity of the Braille alphabet in AlexNet, increasing the sensitivity of the clustering measure. The similar mean dissimilarity in CORnet models, instead, strengthens the similar clustering for Braille and for Latin alphabet stimuli and possibly hints at a tuning to the statistical redundancies with or without lines. The convergence of clustering of Braille and Latin alphabets can be interpreted as evidence of the visual properties being sufficient for the linguistic processing of a stimulus, regardless the presence or absence of line junctions. Alternatively, it can also be seen as influenced by the output class, shared between Braille and Latin alphabet, acting as a lexical entry, with the output layer representing a semantic space. From these results alone, a conclusion is difficult to reach. The representations at the output layer offer a possible clarification in the shortfall of the examined computational models. Primarily, there is a lack of correlation between the patterns found within the networks and a theoretical model of linguistic organization, which orders the different classes of stimuli (Words, Pseudo Words, Non Words, Fake Script) based on the cumulative number of linguistic properties they possess. Even between networks representations, we found mostly within-scripts correlations and not a correlational profile that indicates expertise effects. Both these results do not align with neural data, where alphabets trained converge regardless the visual features (Cerpelloni et al., 2025). Future analyses should look more in depth at the representations within the layers of the networks and at what steps are needed to bridge this gap.

Overall, these results indicate that the visual strategies employed by deep neural networks to process Braille letters do not explain the results found in the behaviour of human participants who learnt Braille(Cerpelloni et al., 2026), nor the neural processing of Braille in expert individuals (Cerpelloni et al., 2025). While further simulations are warranted to understand the nature of this discrepancy, the difference between neuroimaging and computational results suggests a role of additional processes of linguistic computation in the human brain that go beyond the bottom-up vision-based information processing that is implemented in the artificial neural networks that we used. Such additional processes could help human information processing to compensate for the lack of line junctions when having to use Braille as a script, resulting in only minimal delays in learning such script relative to the major delays that we observed in artificial networks. Similarly, interactions with language-related brain areas might help tune visual representations to reflect the linguistic category that stimuli belong to, resulting in a graded selectivity for Real Words, Pseudo-words, Non-words, and Fake script that is absent in the expert visual networks that we tested.

Recently, Chen and colleagues combined human neuroimaging and CLIP models, finding evidence in support of language modulation in vision from the analysis of reduced brain connectivity between VOTC and language areas and the correspondence between CLIP models and brain activity (Chen et al., 2025). This study already points in the same direction of this paper in revealing that, despite their popularity as a model for human vision including reading, bottom-up artificial neural networks do not easily simulate the behavioural and neural findings that we observed with skilled Braille readers. In this context, future models of reading should adapt vision-language models (VLMs; e.g. CLIP or TRIBE v2) to process images of words in addition to images and words or combine visual networks to specific language-processing layers, with the explicit goal of characterising the emergence of linguistic representations from a visual input.

## Author contributions

Conceptualization: FC, OC, HO. Data curation: FC. Formal Analysis: FC. Funding acquisition: OC, HO. Investigation: FC. Methodology: FC, OC, HO. Project administration: OC, HO. Resources: FC, OC, HO. Software: FC. Supervision: OC, HO. Validation: FC. Visualization: FC. Writing – original draft: FC, OC, HO. Author’s contributions are detailed in Figure 5.

**Figure 5.**
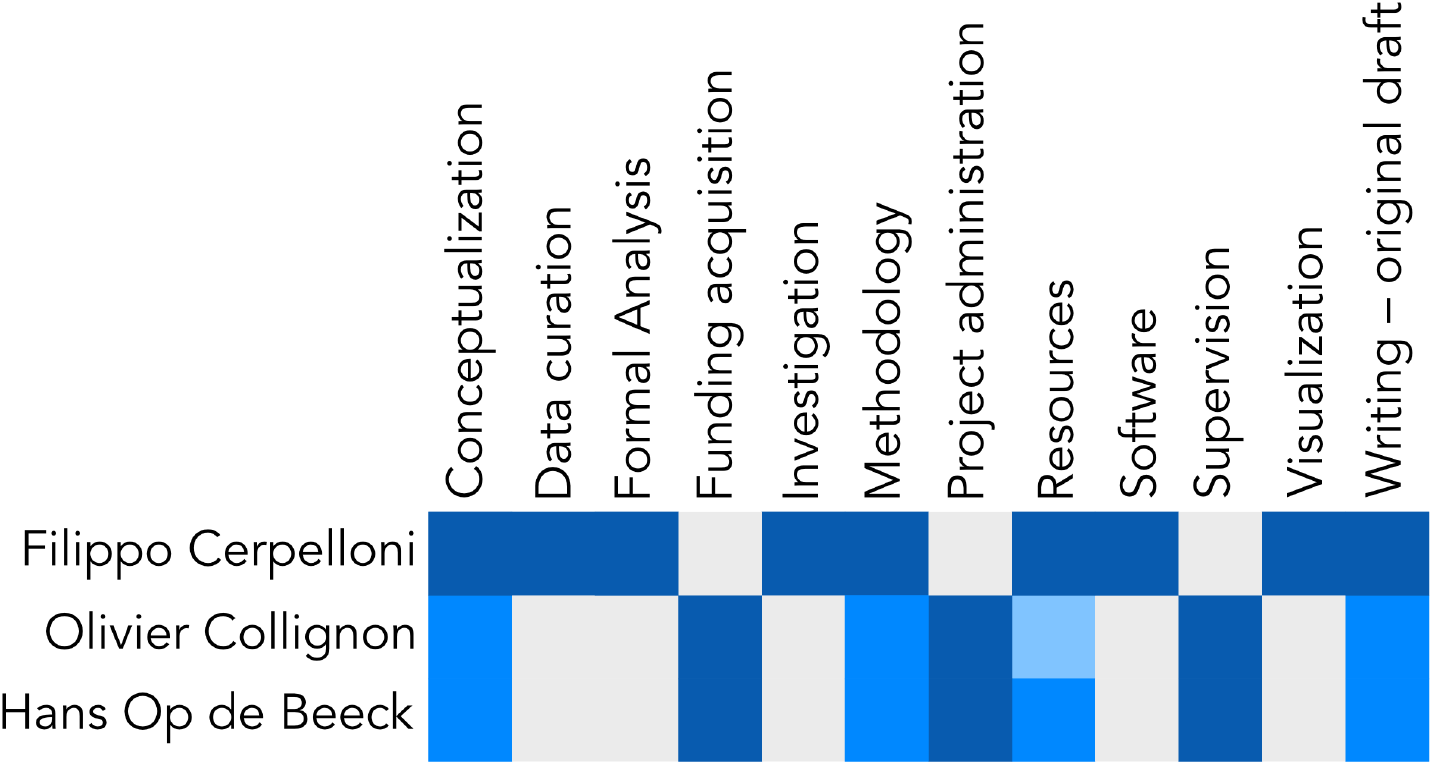
Author contributions. Based on the CRedIT (https://credit.niso.org/) taxonomy. Each contribution was assessed lead’ (dark blue), ‘equal’ (mid blue), ‘support’ (light blue).

## Acknowledgements

The project was funded in parts by a Mandat d’Impulsion Scientifique awarded to OC, a Flagship ERA-NET grant SoundSight (FRS-FNRS PINT-MULTI R.8008.19) awarded to OC, and research project G0D3322N of the Fund for Scientific Research (FWO) Flanders and Methusalem project METH/24/003 awarded to HO. FC received funding from a KU Leuven-UCLouvain cooperation grant.

